# Neutralizing antibody levels and epidemiological information of patients with breakthrough COVID-19 infection in Toyama, Japan

**DOI:** 10.1101/2023.02.27.530346

**Authors:** Hideki Tani, Noriko Inasaki, Takahisa Shimada, Yumiko Saga, Hiroyasu Kaya, Yumiko Maruyama, Sadaya Matano, Hiroyuki Itoh, Tatsuhiko Kashii, Emiko Yamazaki, Shunsuke Yazawa, Masae Itamochi, Kazunori Oishi

## Abstract

Breakthrough infection (BI) after coronavirus disease 2019 (COVID-19) vaccination has exploded owing to the emergence of various SARS-CoV-2 variants and has become a major problem at present. In this study, we analyzed the epidemiological information and possession status of neutralizing antibodies in patients with BI using SARS-CoV-2 pseudotyped viruses (SARS-CoV-2pv). Analysis of 44 specimens diagnosed with COVID-19 after two or more vaccinations showed high inhibition of infection by 90% or more against the Wuhan strain and the Alpha and Delta variants of pseudotyped viruses in 40 specimens. In contrast, almost no neutralizing activity was observed against the Omicron BA.1 variant. Many cases without neutralizing activity or BI were immunosuppressed individuals. The results of this study show that BI occurs even when there are sufficient neutralizing antibodies in the blood due to exposure to close contacts at the time of infection. Thus, even after vaccination, sufficient precautions must be taken to prevent infection.

## Introduction

Over 660 million cases of coronavirus disease 2019 (COVID-19) and over 6.7 million deaths have been confirmed as of January 2023. In response, several vaccines against SARS-CoV-2 have been developed, including mRNA vaccines such as mRNA-1273 of Moderna and BNT162b2 of Pfizer/BioNTech.^1-3^ Vaccination with COVID-19 mRNA vaccines is underway worldwide due to vaccine efficacy in clinical trials. Although the vaccination rate has increased, the number of antibodies has decreased over time, and cases of breakthrough (BT) infection (BI) caused by emerging variants of SARS-CoV-2 have occurred.^4^

In Toyama Prefecture, the Wuhan strain of SARS-CoV-2 was dominant in the early stages, and then the epidemic of the variant changed to Alpha, Delta, and Omicron variants. With the emergence of each variant, the efficacy of mRNA vaccines targeting the Wuhan strain has diminished, and the need for booster vaccine administration has been debated.^5,6^

BI of the COVID-19 vaccine is defined as the detection of SARS-CoV-2 genomic RNA or antigen in respiratory specimens collected from individuals who have received the full recommended dose of the Food and Drug Administration-approved COVID-19 vaccine for at least 14 days.^7^ Several information of infected patients, such as the age, underlying diseases, and vaccine history, as well as the status of possession of neutralizing antibodies at the time of infection, are unknown. Recent reports have shown that individuals with BIs have milder symptoms than those who have not been vaccinated.^8^

In this study, we analyzed the epidemiological information and possession status of neutralizing antibodies in patients with BI using a highly sensitive measurement method with SARS-CoV-2 pseudotyped viruses bearing the spike protein of each variant.^9-11^

## Materials and Methods

### Specimen collection

This study enrolled patients with COVID-19 diagnosed with BI between late July 2021 and early February 2022 at designated medical institutions in Toyama Prefecture. Using the ID numbers of the Toyama Institute of Health (TIH), the specimens were labeled from BT03 to BT51. Several information, including the number of vaccinations, vaccine type, days from vaccination to the onset, date of illness on blood collection, and individual information from the individuals were obtained using a survey form. In addition, sera were collected as early as possible after the diagnosis of COVID-19, and neutralizing activities against viral infection by antibodies in the blood were evaluated.

The severity at the time of blood collection described in the questionnaire was classified as mild, moderate I, moderate II, and severe according to the severity classification of the “New Coronavirus Infectious Disease Treatment Guide Version 7.0”.

### Generation of pseudotyped viruses

Pseudotyped vesicular stomatitis virus (VSV) bearing the SARS-CoV-2 S protein was generated as previously described.^9,12^ The expression plasmids for the truncated S protein of SARS-CoV-2, Wuhan, Alpha, Delta, and Omicron BA.1 were kindly provided by Dr. C. Ono and Prof. Y. Matsuura from the Research Institute for Microbial Diseases, Osaka University. Neutralization of the serum against the pseudotyped viruses was measured as described previously.^9,10^ Briefly, 100-fold dilution of sera in Dulbecco’s Modified Eagle Medium (Nacalai Tesque, Inc., Kyoto, Japan) containing 10% heat-inactivated fetal bovine serum was incubated with each SARS-CoV-2pv or VSVpv at 37°C for 1 h. After incubation, the treated pseudotyped viruses were inoculated into VeroE6/TMPRSS2 cells. The infectivity of the pseudotyped viruses was determined by measuring luciferase activity after 24 h of incubation at 37°C. Samples with each SARS-CoV-2pv or VSVpv treated with healthy donor sera were defined as 100% infection. The neutralization activity in the sera of patients with BI was measured as a reduction in the percentage of infection. If the infection rate was 0.1, 1, or 10%, the neutralization activity was noted as 99.9, 99, or 90%, respectively.

### Statistical analysis

Results are expressed as mean ± standard deviations. Significant differences in the means were determined using the Tukey’s test.

### Ethics approval

This study was performed in accordance with the Declaration of Helsinki and approved by the ethical review board of TIH for the use of human subjects (approval no: R3-2). Written informed consent was obtained from all the patients.

## Results and Discussion

In this study, serum samples were collected from 44 patients with BI. All serum specimens, except for 4 out of 44 cases, were confirmed to have high neutralizing activity that could inhibit the Wuhan strain of SARS-CoV-2pv at a rate of more than 90% (Fig. 1). Blood collection was performed as early as possible from the date of onset to exclude the possible effects of neutralizing antibodies induced after infection. Although nine serum specimens had passed more than 5 days from the date of onset to the date of blood collection, strong neutralizing activities against the pseudotyped viruses were observed. The sera from four specimens showed weak neutralizing activities of no less than 50% against each SARS-CoV-2pv. Three specimens with weak neutralizing activities were collected from a patient receiving immunosuppressive agents (BT22), with HIV infection (BT43), and with malignant lymphoma (BT49). BT43 had acquired immunodeficiency syndrome (AIDS) at the time of diagnosis but had no opportunistic infections and did not take medication; therefore, the CD4 + T cell count (80) was low. BT49 had a stage 4 follicular lymphoma and has received repeated rituximab + CHOP chemotherapy, rituximab + bendamustine, and obinutuzumab + bendamustine therapies. Since all of these patients were immunosuppressed, BI evidently occurred without the production of specific antibodies, even after the administration of the COVID-19 mRNA vaccine. In BT25, the day of illness on the day of blood collection was day 0, and almost no neutralizing activity was detected. Therefore, this case may have also occurred due to the lack of neutralizing antibodies at the time of BI. Next, the neutralizing activity of sera against each SARS-CoV-2pv variant were analyzed. The rate of inhibition in majority of the specimens was highest in the Wuhan strain, followed by the Alpha, Delta, and Omicron variants (Fig. 1). In particular, almost no neutralizing activity against the Omicron variant was observed in the sera of patients with BI.

**Fig. 1.**
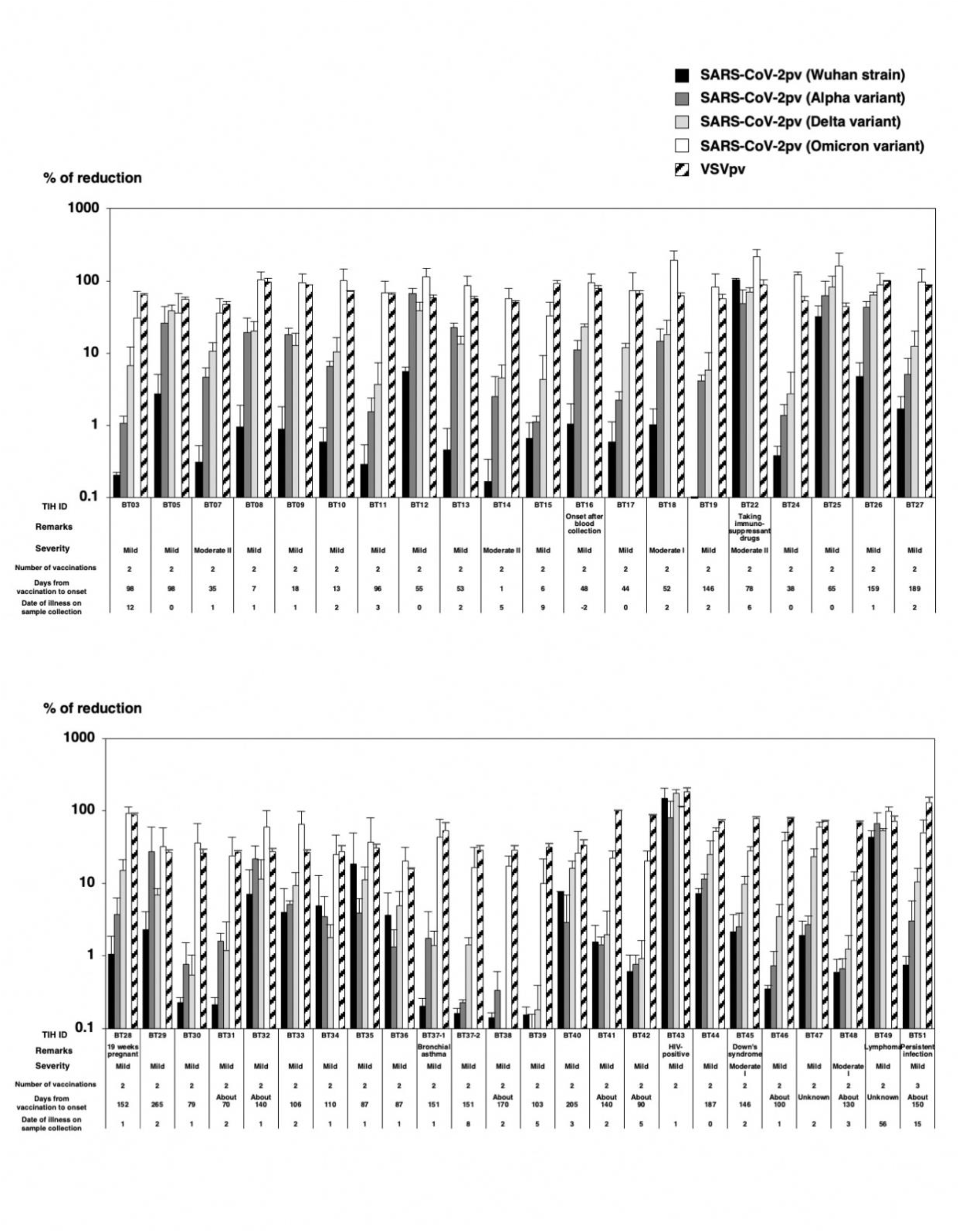
The relative infectivity of SARS-CoV-2pv, Wuhan strain (Wuhan-Hu-1), Alpha variant (B.1.1.7), Delta variant (B.1.617.2), and Omicron variant (BA.1) or VSVpv treated with sera of patients with BI. The medical information of all 44 patients (BT03 to BT51) is shown. Severity was classified into mild, moderate I, and moderate II, according to the severity classification of the “New Coronavirus Infectious Disease Treatment Guide Version 5.0 to 7.0”. The results shown are from three independent assays, with error bars representing standard deviations.

Fig. 2. shows the neutralizing activity of each SARS-CoV-2pv variant according to the period from onset to blood collection. Several patients with BI harbored neutralizing antibodies that could inhibit infection against the Wuhan strain, as well as the Alpha and Delta variants, even in sera collected less than 1 d after the onset of symptoms. Although the neutralizing activity against the Omicron variant was lower than that against the other variants or strain, it increased daily after infection (Fig. 2D). The rate of neutralization increased in all strain or variants 2 days after the onset of symptoms. This may be due to the production of new neutralizing antibodies during infection. However, no significant differences were observed.

**Fig. 2.**
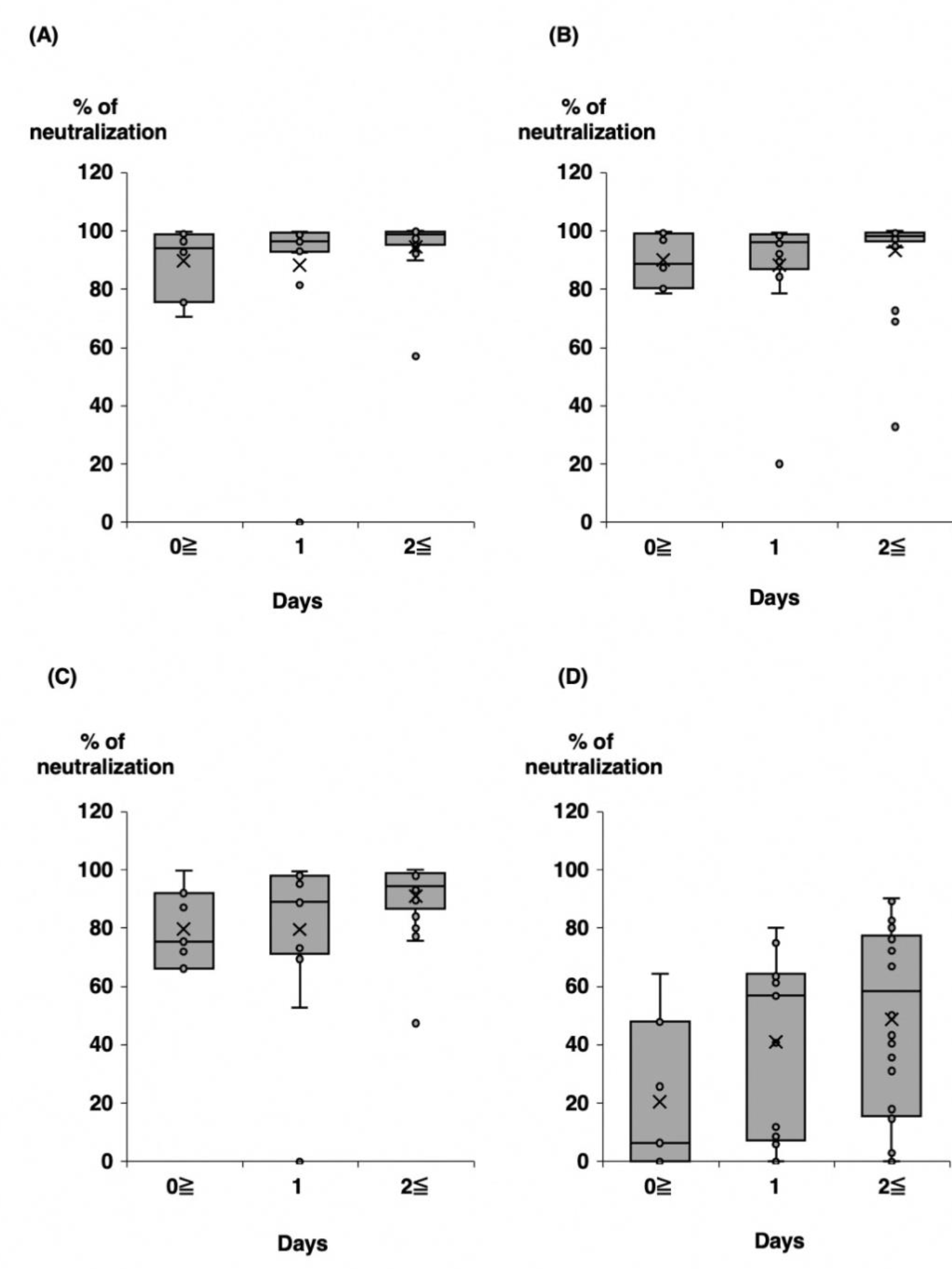
Comparison of the relative neutralizing activity against each SARS-CoV-2pv strain or variant according to the period from the onset to collection of specimens. (A) Wuhan strain, (B) Alpha variant, (C) Delta variant, (D) Omicron variant. ×; mean value, —; median value.

The neutralizing activity against each strain or variant according to the number of days from vaccination to disease onset in patients with BI was also examined (Fig. 3). The neutralizing activity against the Wuhan strain and alpha variant was found to be sufficient within 14 days or even 91 days after vaccination at the time of BI (Fig. 3A and 3B). In contrast, Delta and Omicron variants showed higher neutralizing activity after 91 days than within 14 days of vaccination. This may be due to the increased neutralizing activity due to the booster effect of the infection, as the level of antibodies was not enough against the Delta and Omicron variants. There were also no significant differences in the results of this analysis or the analysis by severity (data not shown).

**Fig. 3.**
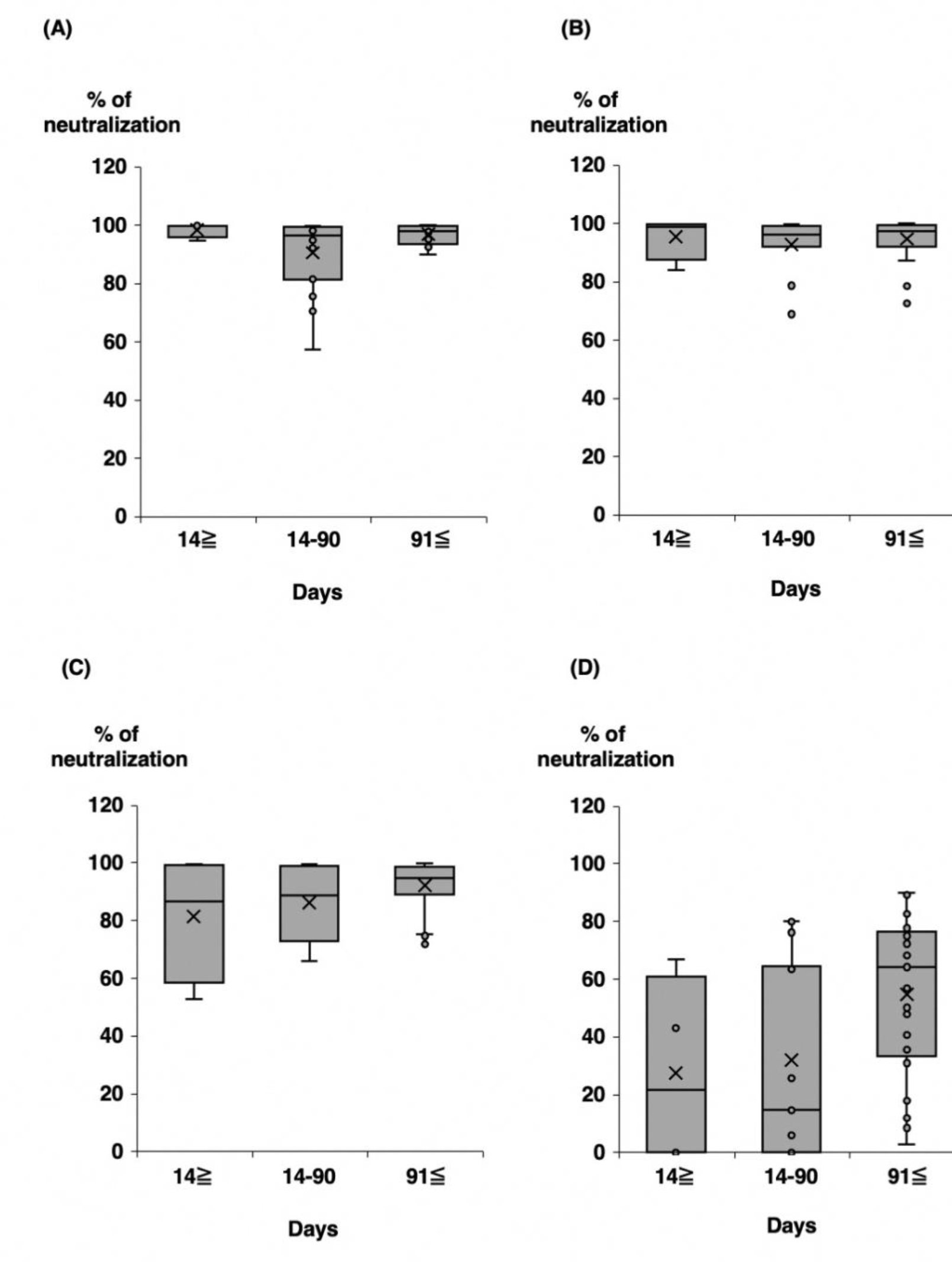
Comparison of the relative neutralizing activity against each SARS-CoV-2pv strain or variant according to period from vaccination to the onset. (A) Wuhan strain, (B) Alpha variant, (C) Delta variant, (D) Omicron variant. ×; mean value, —; median value.

In summary, the present study revealed individual differences in the effectivity of neutralizing antibodies for protection against SARS-CoV-2 infection. BI occurred even in individuals with antibodies possessing a neutralizing activity of 99.9% or higher against the pseudotyped virus. BI is thought to be caused not only by the level of serum neutralizing antibodies but also by exposure to close contacts at the time of infection.

## Author Contribution

Conceptualization and methodology: H.T., N.I., and K.O.; sample collection: H.K., Y.M., S.M., H.I., and T.K.; investigation. N.I., T.S., Y.S., E.Y., S.Y., and M.I.; writing the original draft presentation, H.T.; writing the review and editing, H.T., N.I., and K.O.; supervision, H.T. and K.O.; funding acquisition, H.T. All authors have read and agreed to the published version of the manuscript.

## Acknowledgments

We sincerely thank the Toyama City Public Health Center, Takaoka Welfare Center, Tonami Welfare Center, Niikawa Welfare Center, Chubu Welfare Center, and the Health Division of the Toyama Prefectural Government for the collection, transportation, and arrangement of clinical specimens. We thank Yoko Kanamori and Izumi Kawaguchi for their technical and secretarial assistance.

